# Valuation in major depression is intact and stable in a non-learning environment

**DOI:** 10.1101/074690

**Authors:** Dongil Chung, Kelly Kadlec, Jason A. Aimone, Katherine McCurry, Brooks King-Casas, Pearl H. Chiu

## Abstract

The clinical diagnosis and symptoms of major depressive disorder (MDD) have been closely associated with impairments in reward processing. In particular, various studies have shown blunted neural and behavioral responses to the experience of reward in depression. However, little is known about whether depression affects individuals’ valuation of potential rewards during decision-making, independent from reward experience. To address this question, we used a gambling task and a model-based analytic approach to measure two types of individual sensitivity to reward values in participants with MDD: ‘risk preference,’ indicating how objective values are subjectively perceived, and ‘inverse temperature,’ determining the degree to which subjective value differences between options influence participants’ choices. On both of these measures of value sensitivity, participants with MDD were comparable to non-psychiatric controls. In addition, both risk preference and inverse temperature were stable over four laboratory visits and comparable between the groups at each visit. Neither valuation measure varied with severity of clinical symptoms in MDD. These data suggest intact and stable value processing in MDD during risky decision-making.

## Introduction

Major depressive disorder (MDD) has been associated with impairments in reward processing, and many studies indicate that symptoms of MDD correlate with diminished neural and behavioral responses when rewards are ^1–6^. These studies have typically used reward learning and other tasks that provide feedback about rewards and focused on individuals’ responses at this feedback or ‘reward outcome’ phase (see Rizvi et al.^7^ for review). However, little is known about how depression affects reward valuation during decision-making in the absence of learning and feedback. Understanding whether individuals with MDD have disrupted valuation during decision-making at the ‘decision phase’, separate from reward outcome, will clarify whether individuals with MDD are disrupted overall in reward valuation or more specifically in experiencing rewards. Here, we used a risky decision-making task, a model-based analytic approach, and a repeated measures within-subject design across four visits to investigate whether participants with MDD have intact or disrupted valuation during decision-making in the absence of learning and feedback.

Sixty-nine individuals with current MDD and 41 non-psychiatric controls were recruited for the current study. To investigate ‘value sensitivity’ during decision-making independent from feedback, we asked participants to complete a risky decision-making task (adapted from Holt & Laury^8^ and Dickhaut et al.^9^) (**Fig. 1**). During the task, participants made a series of nine choices between two gambles, one of which was objectively riskier than the other^8^. Each pair of gambles had the same high- and low-payoff probabilities that increased from 10% to 90% in 10% increments along the nine pairs. Participants’ choices between the safer and riskier options, at each payoff and probability combination, were recorded to investigate individual value sensitivity. Participants were paid based on the actual outcome of one of their choices; the outcome was determined after all choices had been made (i.e., no feedback at each decision). This paradigm allowed us to examine valuation during decision-making, independent from potential learning and outcome effects.

**Figure 1.**
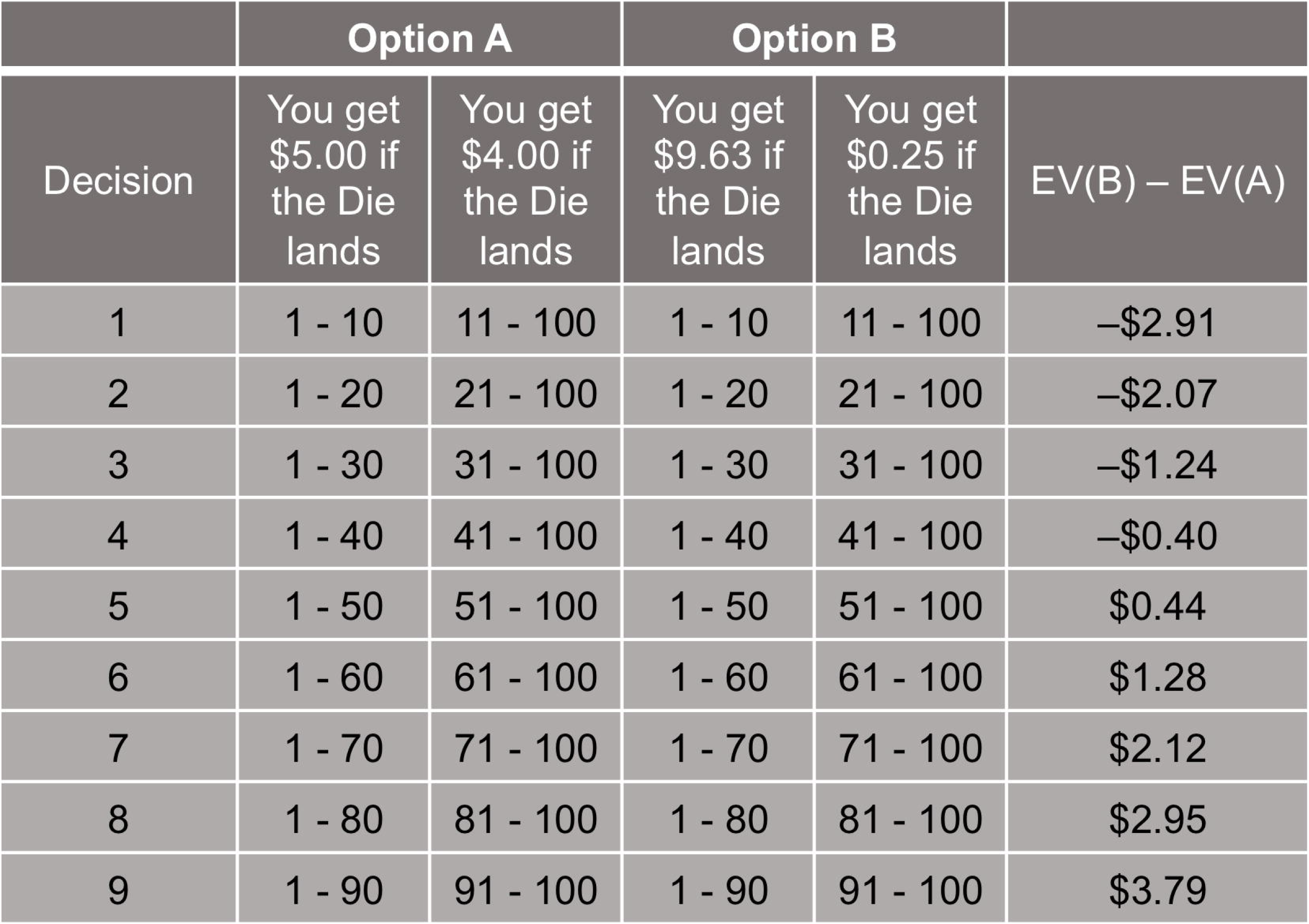
Payoffs and probabilities of paired gambles. Participants played a gambling task that consisted of a menu of probabilities of high and low payoff values. As per Holt & Laury^8^, participants made nine choices between two risky gambles ‘Option A’ and ‘Option B’. The high and low payoffs assigned to each option were fixed as shown here. The probability associated with payoff values was represented as a range of numbers; this allowed participants to easily match the probability of each outcome with a roll of a hundred-sided die; this roll was performed after the task for one randomly selected gamble to determine the final outcome for payoff. The rightmost column shows the expected value differences between the Option A and B. Expected utility theory predicts that a risk neutral individual will choose Option A in decisions 1–4 where EV(B) < EV(A) and Option B in decisions 5–9 where EV(B) > EV(A).

Tasks of this sort are classically used to study individuals’ value-based decision-making under risk^10,11^, and expected utility theory^12^ points to two basic components that account for differences among individuals’ choices in such tasks. The first, ‘risk preference^13,14^ (RP)’ reflects how objective values are subjectively perceived (subjective value) and is quantified by the curvature of a power utility function^12^. The second component determines the degree to which subjective value differences between options affect the probability of choosing one option over the other, and is often referred to as ‘inverse temperature^15^ (IT)’. Both components characterize individual differences in the direction and the degree to which objective values impact individual choices, and thus are used as measures of *value sensitivity* in the current study. Note that each measure explains a different functional relationship between subjective values and decision-making: RP accounts for nonlinear (concave or convex function) subjective valuation and IT is a linear scaling of subjective values (similar to ‘reward sensitivity’ in other MDD studies^1^; see **Methods** for expected utility model specifications). The value sensitivity measures were estimated from individuals’ choices using maximum a posteriori fitting (see **Methods** for parameter estimation procedure).

Participants completed the decision-making task on up to four laboratory visits as part of a longitudinal study; on average, visits were separated by 5.5 weeks (mean of 116.27 days between Time 1 and Time 4 visits). At each visit, participants were instructed that one of their actual choices would be randomly selected and played out to determine their payoff at the end of the visit. The payoff was determined by the values of a gamble selected via random number generator from the participant’s actual choices and a roll of a hundred-sided die (the first determining which gamble would be played and the second determining the payoff). Participants who made less than two visits to the laboratory, had Beck Depression Inventory (BDI-II) scores^16^ > 12 for controls or < 13 at Time 1 for MDD participants, or always chose the option with smaller expected value were excluded from analyses (see **Methods** for numbers of excluded participants for each criterion). The analyzed sample for RP and IT parameter estimation included 33 non-psychiatric controls (14 females; age = 33.00 ± 11.31) and 65 individuals with MDD (48 females; age = 37.92 ± 11.48). See **Table 1** for further demographic information.

**Table 1.** Demographic and symptom data.

|  | Control<br>(*N* = 33) | Major depression<br>(*N* = 65) |
| --- | --- | --- |
| Male/female participants | 14/19 | 17/48 |
| Age (years) | 33.00 ± 11.31 | 37.92 ± 11.48 |
| Verbal intelligence quotient<sup>a</sup> | 105.15 ± 14.90 | 107.06 ± 12.18 |
| BDI-II |  |  |
| Time 1 | 1.55 ± 2.05 (33) | 30.71 ± 7.75 (65) |
| Time 2 | 1.40 ± 2.19 (30) | 22.83 ± 10.70 (53) |
| Time 3 | 1.28 ± 1.79 (29) | 19.12 ± 12.79 (49) |
| Time 4 | 1.64 ± 2.57 (33) | 17.14 ± 13.34 (64) |
| State anxiety |  |  |
| Time 1 | 28.48 ± 6.55 (33) | 49.05 ± 10.82 (65) |
| Time 2 | 27.39 ± 6.35 (31) | 46.47 ± 10.97 (53) |
| Time 3 | 27.86 ± 7.72 (29) | 42.04 ± 12.40 (51) |
| Time 4 | 27.79 ± 6.73 (33) | 39.52 ± 12.76 (64) |
| MASQ subscales |  |  |
| Anhedonic Depression |  |  |
| Time 1 | 45.09 ± 8.97 (33) | 83.11 ± 9.12 (65) |
| Time 2 | 43.29 ± 9.84 (31) | 71.63 ± 14.39 (52) |
| Time 3 | 42.86 ± 10.72 (29) | 66.47 ± 18.96 (51) |
| Time 4 | 43.38 ± 10.36 (32) | 65.00 ± 16.86 (65) |
| Anxious Arousal |  |  |
| Time 1 | 18.55 ± 1.99 (33) | 26.85 ± 7.25 (65) |
| Time 2 | 18.39 ± 1.87 (31) | 23.79 ± 6.90 (52) |
| Time 3 | 18.69 ± 3.29 (29) | 23.80 ± 8.31 (51) |
| Time 4 | 18.28 ± 1.49 (32) | 23.62 ± 8.21 (65) |
| GD: Anxiety |  |  |
| Time 1 | 14.48 ± 2.59 (33) | 25.03 ± 6.97 (65) |
| Time 2 | 13.42 ± 2.47 (31) | 20.81 ± 6.18 (52) |
| Time 3 | 14.00 ± 2.60 (29) | 20.59 ± 7.57 (51) |
| Time 4 | 13.44 ± 2.06 (32) | 18.95 ± 6.86 (65) |
| GD: Depression |  |  |
| Time 1 | 15.72 ± 2.82 (33) | 38.94 ± 8.91 (65) |
| Time 2 | 15.16 ± 2.27 (31) | 31.10 ± 9.57 (52) |
| Time 3 | 15.07 ± 3.62 (29) | 27.49 ± 11.08 (51) |
| Time 4 | 15.09 ± 2.44 (32) | 26.68 ± 11.74 (65) |
| GD: Mixed |  |  |
| Time 1 | 22.55 ± 4.49 (33) | 45.91 ± 8.41 (65) |
| Time 2 | 21.03 ± 4.53 (31) | 39.08 ± 9.13 (52) |
| Time 3 | 20.79 ± 3.80 (29) | 37.00 ± 11.01 (51) |
| Time 4 | 21.38 ± 4.83 (32) | 34.55 ± 11.16 (65) |

BDI-II, Beck Depression Inventory, Second Edition; State anxiety, Spielberger's State Anxiety Inventory; MASQ, Mood and Anxiety Symptom Questionnaire; GD, General Distress; <sup>a</sup>Verbal intelligence quotient scores were measured with the Wechsler Test of Adult Reading (WTAR). Numbers of participants who were included for calculating mean and standard deviation of each questionnaire score are noted in parentheses; see **Supplementary Fig. S1** for the BDI-II, SAI, and MASQ scores in the smallest subset of included participants (visiting all four times), and see **Methods** for details about inclusion criteria.

## Results

### Valuation is comparable between MDD and non-psychiatric control participants

To compare value sensitivity in MDD participants with that of non-psychiatric controls, we estimated each individual’s risk preference and inverse temperature for each visit, and first compared the means of these parameters between groups (see **Methods** for details about parameter estimation). Thus, RP and IT at each of the four visits were computed for each individual, for participants who visited all four times (*N*_control_ = 28, *N*_MDD_ = 47). Group mean parameter values were: RP_control_ = 0.50 ± 0.31; RP_MDD_ = 0.46 ± 0.31; IT_control_ = 3.41 ± 0.41; and IT_MDD_ = 3.25 ± 0.43 (mean ± s.d.). Note that both the MDD and non-psychiatric control groups showed risk aversion (RP < 1) consistent with Holt & Laury^8^. Across four laboratory visits, participants with MDD showed comparable RP and IT to that of non-psychiatric controls (**Fig. 2ai, 2bi**; RP: F(1, 219) = 0.63, *P* = 0.43; IT: F(1, 219) = 2.68, *P* = 0.11; Group × Time mixed-design ANOVAs with rank transformation^17^). These results indicate that MDD and non-psychiatric control participants have comparable linear and nonlinear value sensitivities during decision-making.

**Figure 2.**
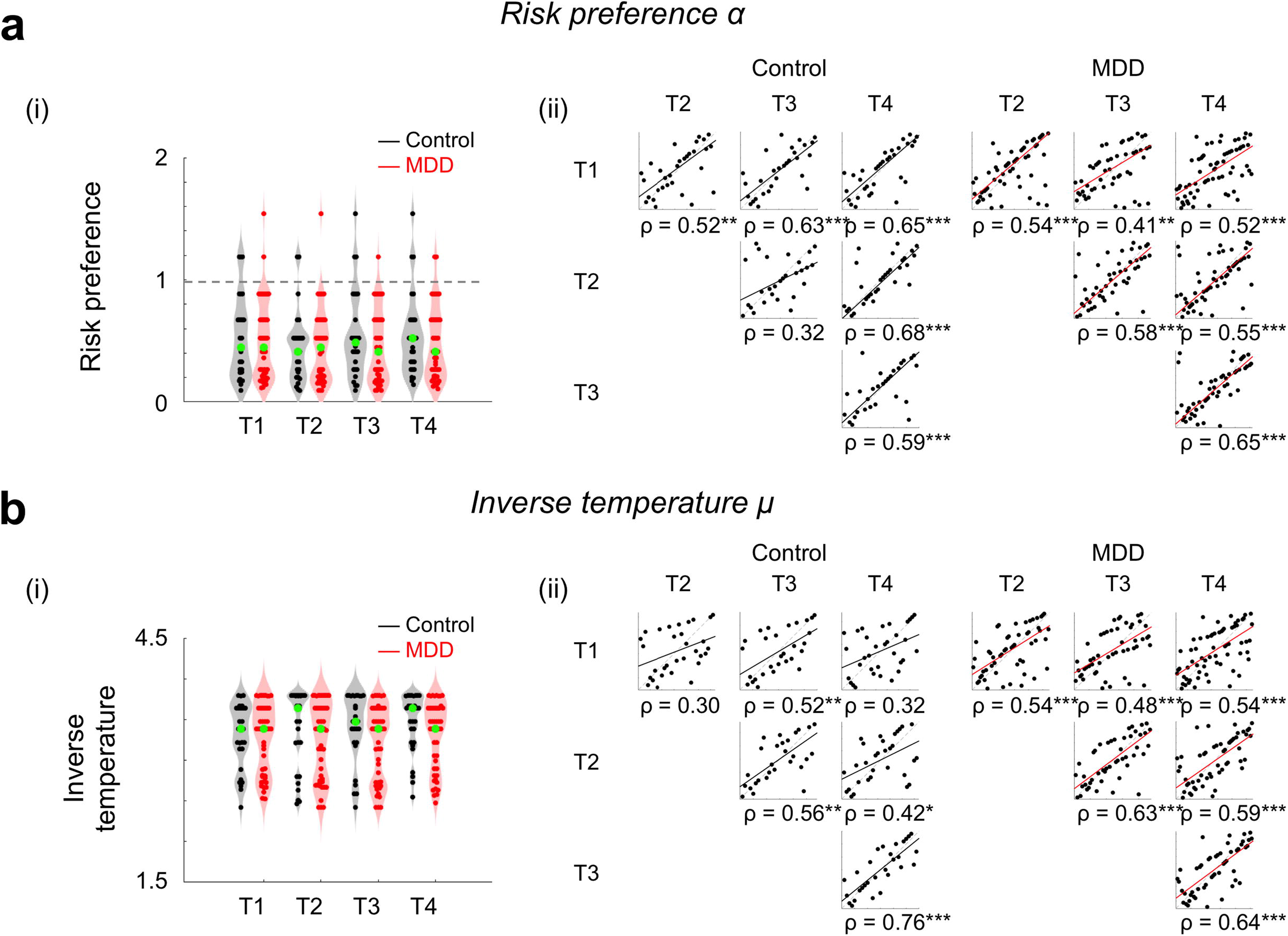
Estimated value sensitivities are comparable between non-psychiatric controls and individuals with MDD, and stable across visits. We used a standard power utility function and softmax choice rule to identify separate ‘risk preference’ and ‘inverse temperature’ parameters to explain nonlinear and linear value sensitivities in decision-making. **(ai, bi)** Estimated RP and IT were stable across four repeated visits for both MDD and control participants. Across the repeated visits, both RP and IT were comparable between the control and MDD groups (no main effect of group using mixed-design ANOVA with rank transformation). The gray dotted line indicates risk neutrality (RP = 1). Each point represents an individual participant; group medians are indicated in green. Gray and red shades show the distribution of data points along the y-axis. **(aii, bii)** Spearman’s correlation coefficients were calculated to test whether the rank order of the parameters among individuals was consistent between visits to the lab (([1^st^ vs 2^nd^ visit], [1^st^ vs 3^rd^ visit], … [3^rd^ vs 4^th^ visit]). See **Supplementary Table S1** for statistical results. Each point represents an individual participant, and the color-coded lines are the robust regression line between measures from two visits. The x- and y-axes each represent the rank order of individual participants at each visit (for simplicity, not labeled here); \**P* < 0.05, \*\**P* < 0.01, \*\*\**P* < 0.001, uncorrected; all correlations were significant after applying multiple comparison correction (FDR *q* < 0.0001).

### Valuation is stable over time for both MDD and non-psychiatric control participants

Previous studies have shown that risk preferences measured with variations of the Holt & Laury task^8^ are stable over time in unselected control individuals, particularly when model-based estimates were used^18^,^19^. As an initial test of RP and IT stability within MDD and control participants, we compared within group means over time; these analyses indicate that neither parameter differed over time for either group (RP_control_: **χ**^2^(3, 81) = 2.12, *P* = 0.55; RP_MDD_: **χ**^**2**^(3, 138) = 0.66, *P* = 0.88; IT_control_: **χ**^**2**^(3, 81) = 2.94, *P* = 0.40; IT_MDD_: **χ**^**2**^(3, 138) = 2.40, *P* = 0.49; Friedman’s tests, analogous to non-parametric repeated measures ANOVAs; only participants who visited all four times were examined here (*N*_control_ = 28, *N*_MDD_ = 47)). Adopting the approach of previous studies for measuring temporal stability, we also examined the stability of RP and IT within controls and participants with MDD by correlating the value of each parameter between pairs of visits ([1^st^ vs 2^nd^ visit], [1^st^ vs 3^rd^ visit], … [3^rd^ vs 4^th^ visit]) (**Fig. 2aii, 2bii**). Both control and MDD participants showed moderate to high stability in both RP and IT, respectively (mean correlation coefficients: RP_control_: Spearman ρ = 0.57; RP_MDD_: ρ = 0.54; IT_control_: ρ = 0.48; IT_MDD_: ρ = 0.57; see **Fig. 2aii** and **2bii** for full correlation matrix; **Supplementary Table S1** for sample size at each visit). Note that the proportion of risky choices, a model-free measure of risk preference, was also stable over time in both the MDD and control groups (see **Supplementary Fig. S2** for model-free risk preference stability over time). These significant correlations indicate that for MDD and control participants, the risk preference and inverse temperature measures of value sensitivity at the decision phase are stable over time.

### Valuation does not vary with severity of clinical symptoms in MDD

Given previous reports that reward sensitivity at decision outcome varies with symptoms in depression^1,20–22^, we also examined whether RP and IT varied systematically with depressive or anxious symptoms. Symptoms were measured using the BDI-II^16^, Spielberger’s State Anxiety Inventory (SAI)^23^, and the five subscales of the Mood and Anxiety Symptoms Questionnaire (MASQ; Anhedonic Depression, Anxious Arousal, General distress (GD):Anxiety, GD:Depression, and GD:Mixed)^24^; correlations were performed within the MDD group. None of the clinical symptom scores or changes in symptoms over time were related to MDD participants’ RP or IT parameter values (see **Supplementary Fig. S1**, **Supplementary Table S2,** and **S3** for statistical test results). These data demonstrate that individual differences in value sensitivity during decision-making are not explained by clinical characteristics of MDD.

## Discussion

The current study used a risky decision-making task to investigate MDD individuals’ value sensitivity at the decision phase independent from learning and feedback. The within subjects repeated-measures design allowed us to examine the stability of the value sensitivity measures, and the model-based approach dissociated linear (inverse temperature) and nonlinear (risk preference) value sensitivities that together determine behavioral choices during risky decision-making.

A few previous studies have used risky decision-making paradigms and measured MDD individuals’ risk preferences. The results, however, have been inconsistent. Some studies reported decreased risk seeking behavior in individuals with MDD^20,25,26^, while other studies reported comparable risk preferences between individuals with MDD and healthy individuals^27,28^. In the current study, we showed that risk preferences (nonlinear value sensitivity) in individuals with MDD are comparable with those of healthy individuals. The stability of risk preferences was tested across four repeated visits, and consistent with previous findings in unselected control individuals^18,19,29^, MDD participants showed stable risk preferences over time (c.f., model-free measures showing lower reliability^30–32)^. In addition to estimating risk preference, we examined inverse temperature (linear value sensitivity, similar to ‘reward sensitivity’ in other MDD studies^1^) at the decision phase, and showed that MDD participants have stable and comparable inverse temperature compared with non-psychiatric controls. In addition, none of the clinical symptom severity measures within participants with MDD were related to individual differences in risk preference or inverse temperature. These results indicate that in contrast with previous decision-making studies showing blunted valuation at the outcome phase in MDD^1^, neither linear nor nonlinear value sensitivity at the decision phase in MDD was different from that of controls.

To date, studies examining valuation in MDD have primarily focused on the outcome phase of reward learning tasks and shown impaired valuation, including diminished neural reward responses^33–35^, reduced learning rate^36^, lower reward sensitivity^1^, or enhanced exploration (more frequent choice shifting)^37,38^ in participants with MDD. A few other studies have used various non-learning tasks and have suggested that individuals with MDD have low motivation for monetary reward^20,39,40^; however, in these studies, the focus was also on responses at the outcome phase^21,22^. Unlike the abundant literature about responses to reward outcome (particularly during reward learning), little is known about whether individuals with MDD have intact ability to process and compare values during decision-making when no learning is required. The current study provided no outcome feedback during the task and thus focused on the decision phase dissociated from learning and reward experience. These data showed that during the decision phase, participants with MDD have value processes comparable to that of healthy individuals. This is consistent with previous studies showing intact neural responses in individuals with MDD during reward anticipation (prior to outcome)^41,42^. Together, the present data indicate that individuals with MDD have intact valuation when reward contingencies are fully known (no reward learning required) and suggest that previously reported valuation deficits in MDD are specific to the outcome phase of tasks in which rewards are experienced and learning occurs.

In MDD, intact valuation, dissociated from learning, may provide mechanistic insight about behavioral activation therapies for depression^43^. These type of therapies engage individuals with potential positive reinforcers (rewards) in a structured manner and, in doing so, allow individuals with MDD to largely bypass disrupted learning processes. That is, behavioral activation provides a guided learning environment wherein engagement and experience of action-reward contingencies are enforced, allowing for the value of rewards to evolve from being unsampled and ambiguous to sampled and fully known. Once these values are known, intact decision processes such as those identified here allow individuals to engage in healthy choices. As our data indicate, when action-reward contingencies are fully known, participants with MDD show intact valuation during decision-making. We speculate that this state is comparable to the endpoint of successful behavioral activation wherein the experience of reward is restored. In brief summary, the current study suggests specificity of previously reported value processing disruptions in MDD, informs the conditions under which sensitivity to reward values is preserved, and offers the possibility that learning about reward values, rather than discriminating among values when making decisions, may be a mechanistic target for intervention in MDD.

## Methods

### Participants

Fifty non-psychiatric controls and 80 individuals with MDD were recruited as part of a larger ongoing study examining neural substrates of treatment response in MDD (neural and treatment data will be analyzed as part of another manuscript). Among these participants, we included individuals who at least participated in both Time 1 and 4 laboratory visits to maximize the time interval for test-retest reliability. These inclusion criteria yielded 41 non-psychiatric controls and 69 individuals with MDD for the present study. Basic inclusion/exclusion criteria were initially assessed via telephone and were confirmed during the first laboratory visit using the Structured Clinical Interview for DSM-IV-TR Axis I Disorders – Research Version – Patient Edition (With Psychotic Screen) (SCID-I/P)^44^ and selected modules of the Mini-International Neuropsychiatric Interview (M.I.N.I.)^45^. At study intake, individuals in the MDD group met DSM-IV criteria for MDD and/or dysthymia while individuals in the control group did not meet criteria for any current Axis I disorder. Exclusion criteria for all participants included contraindications to magnetic resonance imaging (MRI) and history of neurological disease. Following the initial screening visit (Time 1), participants returned to the lab up to three times; on average, there were four-week intervals in between each visit. All participants provided written informed consent and were given instruction about the task. The study was approved by the Institutional Review Board of Virginia Tech. Three controls whose BDI-II scores were above the non-depressive range (i.e., greater than 12) at any visit and two individuals with MDD who had BDI-II scores in the non-depressive range (i.e., less than 13) at Time 1 were additionally excluded from analyses^46^. Five controls and two individuals with MDD who always chose the option with smaller expected value were also excluded. Therefore, the analyzed sample for RP and IT parameter estimation included 33 healthy controls (14 females; age = 33.00 ± 11.31) and 65 participants with MDD (48 females; age = 37.92 ± 11.48). See **Table 1** for additional demographic information.

### Experimental procedure

Participants made a series of nine choices between two gambles, one of which was objectively riskier than the other (adapted from Holt & Laury^8^) (**Fig. 1**). Each pair of gambles had the same high- and low-payoff probabilities that varied from 10% to 90% in 10% increments along the nine pairs. Payoff spreads between high- and low- payoffs were fixed for each option; ‘Option A’ had $5.00 and $4.00, and ‘Option B’ had $9.63 and $0.25 as potential payoffs. Participants were paid based on the actual outcome of one of their choices; the payoff was determined by the values of a gamble selected via random number generator from the participant’s actual choices and a roll of a hundred-sided die (the first determining which gamble among those presented in multiple tasks included for the larger ongoing study, would be played and the second determining the payoff).

### Model-free analyses

For model-free behavioral analyses, the proportion of choices of the risky option (P(risky)) among the nine pairs of gambles was used as a measure of risk preference. Given the expected value (EV) between pairs of choices (**Fig. 1**), a risk neutral individual should show P(risky) = 5/9 ≈ 0.56 (as per expected utility theory, a risk neutral individual is expected to choose Option B in the trials where EV(B) > EV(A), decisions 5-9, and to choose Option A in the trials where EV(B) < EV(A)). Higher P(risky) thus indicates risk seeking. P(risky) was calculated per visit and used to examine stability of model-free risk preferences over time in each group.

### Estimates of individual risk preference

We applied expected utility theory^12^ to estimate each individual’s risk preference (RP) and inverse temperature (IT) that predict the individual’s choices. We used a standard power utility function and softmax choice rule as described below:

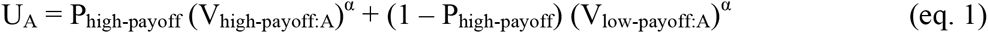

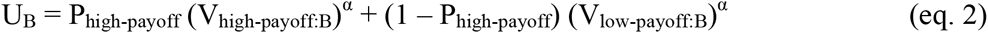

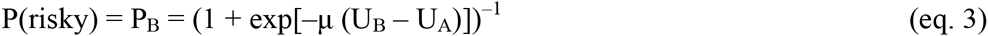

where U_A_ (U_B_) is the utility of the Option A (Option B), P is the probability of earning a payoff, V represents the payoff amount for each gamble, α is the *risk preference,* and µ is the *inverse temperature.* The estimated RP, α, indicates whether an individual is risk averse (0 < α < 1), risk neutral (α = 1), or risk seeking (α > 1). The estimated IT, µ, indicates how sensitive an individual is to the utility differences between the two gambles; larger µ indicates higher sensitivity to utility differences and µ ≈ 0 indicates utility (subjective value) insensitivity.

To achieve a more stable parameter estimation for each individual, we adopted a hierarchical model structure of the population^47^ in which it is assumed that a participant *i*'s parameters (µ_*i*_ and α_*i*_) are sampled from the population’s parameter distribution. Of importance, both controls and participants with MDD were considered to share the same group-level (the population) distribution (equal prior), which allowed us to compare the two participant groups in the further analyses. This is a conservative approach, because the equal prior does not introduce potential bias about different parameter distributions between groups. Based on these assumptions, we estimated the group-level parameter distribution for each parameter and set the distribution as a prior for individual estimation (maximum a posteriori (MAP) estimation). In the current study, we set the group-level distribution of each parameter as a gamma distribution^48^ with a shape parameter, k, and a scale parameter, θ, (µ *~* Γ(k_µ_, θ_µ_); and α ~ Γ(k_α_, θ_α_)). For each iteration of the group-parameter estimation (max iteration of 15,000), 100 random samples were drawn from each parameter distribution for each participant, and the average of the calculated likelihoods were used as an approximation of the integral in the following equation:

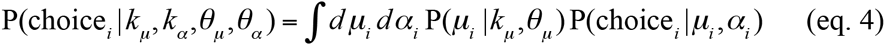

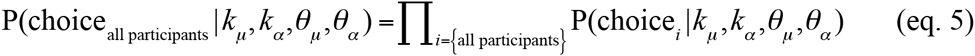

Note that all participants visited at least twice, including the 1st and the 4th visits.

To allow for RP and IT values to vary across time independent from subject level information, we did not provide any information about subjects’ identity in the estimation step; that is, behavioral choices from a participant’s 1st and 4th visits were considered decision patterns from two independent participants. Note that estimated value sensitivities for the same subject from repeated visits were considered as repeated-measures for post estimation stability testing. To apply this method, we used 196 sets of behavioral choices for the group-level parameter estimation ([33 HC + 65 MDD] × [1st visit + 4th visit]; only 1st and 4th visits were used to provide an equal amount of choice information from each individual participant). The group-level parameters were used to define each parameter’s prior distribution for individual-level estimation, which was equally applied to individual-level estimations for all four visits. We fit the data using MAP, with posterior function as below.

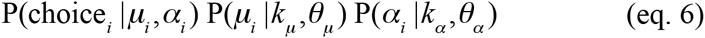

All parameter estimations were conducted with custom MATLAB R2015b (MathWorks) scripts and the fminsearch function in MATLAB with multiple initial values.

### Clinical measures

At each visit, participants completed a battery of self-report measures to assess current depression and anxiety symptoms. Depressive symptom severity was measured using the BDI-II and the Anhedonic Depression subscale score of the MASQ. Anxiety symptom severity was measured using the state scale of the SAI (Spielberger’s Anxiety Inventory) and the Anxious Arousal subscale of the MASQ. Additionally, general distress (GD) related to depressive symptoms, anxious symptoms, or a mixture of the two were measured using the MASQ subscales, GD: Anxiety, GD: Depression, and GD: Mixed, respectively.

### Statistical analyses

We examined if model-free risk preference (proportion risky choices) and model-based measures of value sensitivity (inverse temperature and risk preference) were consistent across multiple visits. IT and RP measures in both participant groups were not normally distributed (Shapiro-Wilk test *P* < 0.01 for IT and RP in each group and in each visit), and thus non-parametric tests were used as appropriate and available. First, to compare the means of IT and RP across four laboratory visits and between groups, we used mixed-design ANOVA where visit number (Time 1, Time 2, Time 3, Time 4) was the within-subject factor and diagnostic group (MDD, control) was the between-subject factor. Parameters were first rank transformed and then inserted for mixed-design ANOVA^17^. In addition, we used Friedman’s test to examine whether IT and RP across four visits were stable or not, within each group. Second, Spearman’s correlations between risk preference measures from two different visits (‘1st visit’ (T1) vs T2, T1 vs T3, T1 vs T4, T2 vs T3, T2 vs T4, and T3 vs T4) were calculated to test if the rank-order of risk preference within each group is consistent across multiple visits. All statistical tests were two-sided. False discovery rate (FDR) adjusted *q*-values where indicated were reported for multiple comparisons^49^. MATLAB R2015b was used for all statistical tests.

## Acknowledgements

We thank J. Lee, R. McNamara, and C. Rosoff for their research support and gratefully acknowledge discussions with A. Solway, V. Brown, and S. Ball. The work was supported in part by the US National Institutes of Health (MH091872, MH087692, MH106756 to P.H.C; DA036017 to B.K.-C.).

## Author Contributions

J.A.A., B.K.-C., and P.H.C. designed the experiments; D.C. and K.K. analyzed the data; D.C., K.K., J.A.A., K.M, B.K.-C., and P.H.C. discussed the analyses and results; D.C. and P.H.C. drafted the manuscript; D.C., K.K., J.A.A., K.M, B.K.-C., and P.H.C. revised and approved the submission.

